# The Role of Objecthood and Animacy in Apparent Movement Processing

**DOI:** 10.1101/2022.08.22.504783

**Authors:** Emiel Cracco, Tilia Linthout, Guido Orgs

**Affiliations:** Department of Experimental Clinical and Health Psychology, Ghent University, Belgium; Department of Psychology, Goldsmiths, University of London, London, UK; Department of Music, Max Planck Institute for Empirical Aesthetics, Frankfurt, Germany

## Abstract

Movement perception involves both motion and form processing. Previous research has shown that processing along the motion pathway requires a familiar but not necessarily a human shape. However, the role of objecthood and animacy in the form pathway is less clear. Here, we used EEG frequency tagging to study how objecthood and animacy influence both posture processing and the integration of postures into movements. Specifically, we measured brain responses to repeating sequences of well-defined or pixelated images (objecthood) of human or corkscrew agents (animacy) performing fluent or non-fluent movements (movement fluency). The results revealed that movement processing was sensitive to objecthood but not to animacy, whereas posture processing was sensitive to both. Thus, our results indicate that movement processing in the form pathway requires a familiar shape, but not necessarily a human shape. Instead, stimulus animacy appears to be relevant only for posture processing.

## Introduction

Processing other people’s movements is imperative for adaptive social behavior (Pavlova, 2012). Models of biological motion perception argue that there are at least two pathways involved in this process (Giese & Poggio, 2003; Lange & Lappe, 2006), a *motion* and a *form* pathway. In the motion pathway, biological motion is processed in terms of movement kinematics and constraints (e.g., Grossman & Blake, 2001; Troje, 2013). In contrast, in the form pathway, biological motion is processed as a sequence of specific limb configurations (i.e., body postures; e.g., Orgs et al., 2011; Shiffrar & Freyd, 1990). Evidence for a form pathway comes from apparent motion studies, showing that sequences of two or more static images induce a vivid motion percept as long as the interval between images is consistent with the duration of the implied movement (Grosjean et al., 2007; Shiffrar & Freyd, 1990).

Motion and form processing of biological movement have shown to rely on distinct neural mechanisms. Indeed, whereas motion information is primarily processed in motion-specific areas such as the middle temporal cortex (Mather et al., 2016; Vaina et al., 2001), form information is processed mainly in body-specific areas such as the extrastriate body area (Orgs et al., 2016; Vaina et al., 2001), though both types of information eventually converge in higher-order areas like the superior temporal sulcus (Pitcher & Ungerleider, 2021; Vaina et al., 2001). What is less clear, however, is the extent to which biological motion processing along the motion and form pathways can be triggered by shapes other than human bodies. In two studies, Jastorff et al. (2006, 2009) used point-light stimuli, which contain minimal form information, to investigate this question in the motion pathway. The results revealed that, after training, human and non-human stimuli were processed in the same way, as long as they had a clear underlying shape (as opposed to scrambled motion). This suggests that movement processing in the motion pathway is not specific to human body stimuli, but rather reflects a more general mechanism that integrates local motion cues into a global movement pattern consistent with a learned, familiar shape (Giese & Poggio, 2003; Lange & Lappe, 2006).

Similar to Jastorff et al. (2006, 2009), research suggests that movement processing along the form pathway is likewise disrupted by manipulations that affect stimulus shape, such as pixelation (Orgs et al., 2011, 2016). However, whether it can also proceed without a human shape is not yet known. On the one hand, a human shape may be necessary because the form pathway involves brain areas known to display human specificity (Downing, 2001; Orgs et al., 2016). On the other hand, recent work suggests that even in those brain areas, animacy is not represented categorically, but on a continuum from more to less alive (Sha et al., 2015; Thorat et al., 2019). Here, we used a recently developed frequency tagging task to directly test if apparent movement perception is specific to human shapes or generalizes to non-human shapes as well (Cracco, Lee, et al., 2022). In this task, a sequence of 12 images is shown repeatedly using a fixed image presentation rate (Figure 1). Importantly, the sequence is symmetric, with the second half of the images mirroring the first half. As a result, three brain responses are elicited: a response coupled to the frequency of image presentation (base rate), a response coupled to the turning point in the image sequence (half cycle rate), and a response coupled to the completion of the entire sequence, that is image repetition (full cycle rate).

**Figure 1.**
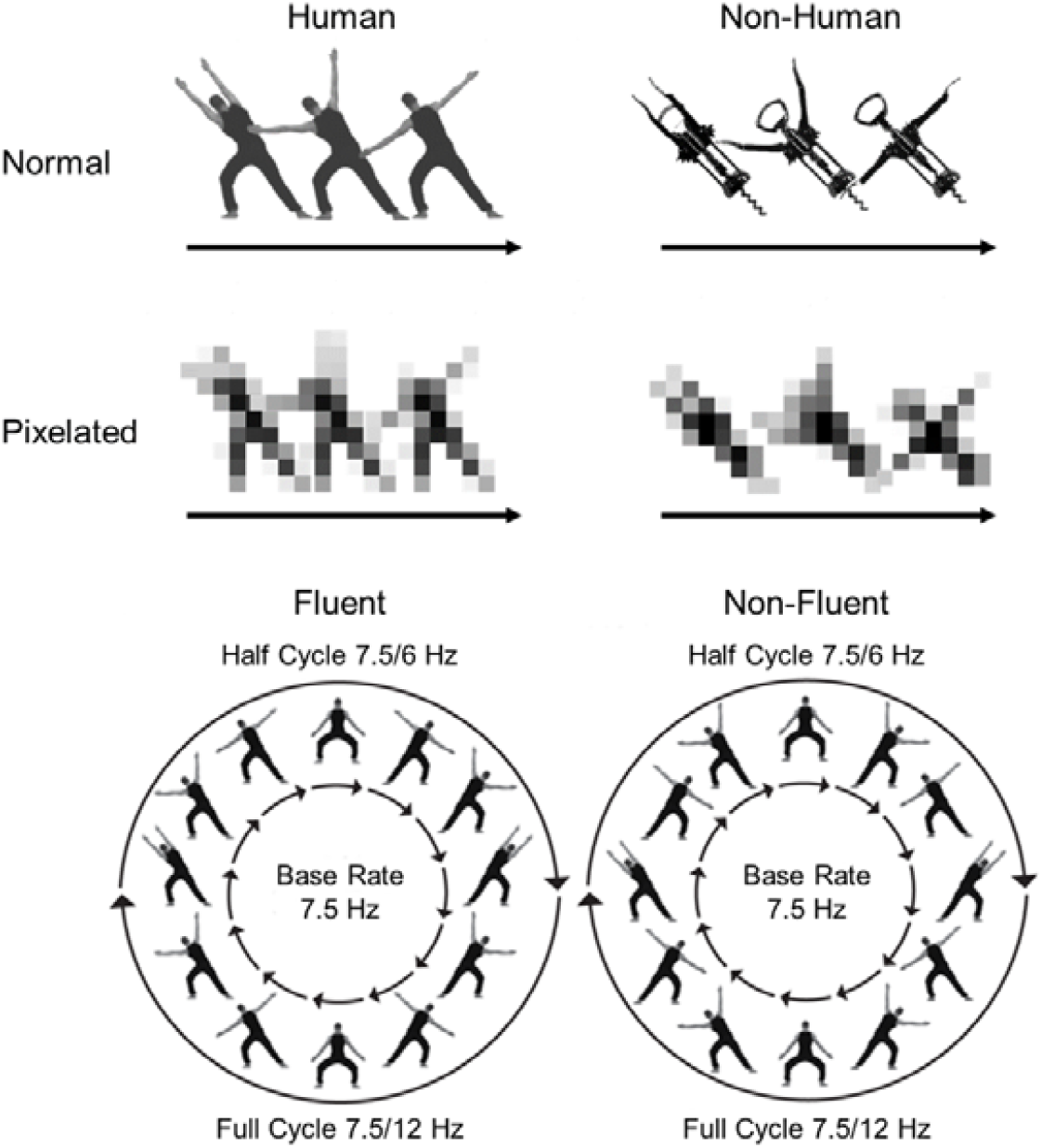
Stimuli and paradigm. The top shows the four types of stimuli used in the study. The bottom shows the structure of the fluent and non-fluent sequences. The latter figure is adapted from Figure 2 of Cracco, Lee, et al. (2022), where it was published under a CC BY license.

To study apparent movement processing, Cracco, Lee, et al. (2022) compared two types of sequences. In fluent sequences, images were ordered to produce the percept of a human agent making fluent movements. In non-fluent sequences, they were instead ordered to disrupt this percept. Crucially, movements in the fluent condition were executed at half cycle rate. This made the half cycle point more salient and led to stronger half cycle responses, thereby capturing movement processing (Cracco, Lee, et al., 2022). In contrast, full cycle and base rate responses both showed the opposite pattern, with stronger responses for non-fluent than for fluent sequences. In line with evidence that scrambling the image order of an apparent movement sequence causes image processing to take over (e.g., Downing et al., 2006), these results thus indicate that non-fluent sequences were processed as *image sequences*, at base rate (presentation of images) and full cycle rate (completion of an image sequence), whereas fluent sequences were processed as *movement sequences*, at half cycle rate.

Cracco, Lee, et al. (2022) found that half cycle amplitudes were weaker for inverted than for upright body stimuli, suggesting that the processing of apparent movement is orientation-specific (see also Orgs et al., 2011). However, whether it is also specific to human stimuli is not yet known. To investigate this question, we manipulated two image features: objecthood (normal vs. pixelated) and animacy (human vs. corkscrew; Figure 1). In line with the corresponding research on biological movement processing in the motion pathway (Jastorff et al., 2006, 2009), this allows us to disentangle the role of both these features. Indeed, even though a corkscrew is clearly not alive (animacy), it still has a familiar and well-defined shape (objecthood) that moves similar to a human body, its “arms” rotating across a central “torso”. If movement processing in the form pathway is influenced not just by objecthood (e.g., Orgs et al., 2011, 2016) but also by animacy, we should find specific effects of both features on the half cycle amplitudes. In particular, we should find an interaction between pixelation and animacy, such that half cycle responses are strongest for sequences showing non-pixelated human images. Moreover, if these effects are specific to movement processing, we should further find that they are stronger for fluent than for non-fluent sequences, given that only fluent sequences produce a movement percept (Cracco, Lee, et al., 2022).

### Open Science Statement

This study was preregistered: https://aspredicted.org/8RH_NXF. Deviations from the preregistration are mentioned and justified in the methods. An example video, the preprocessed data, and the analysis script will be made available on the Open Science Framework.

## Methods

### Participants

Because this is the first study to compare apparent movement processing of human and non-human agents, we did not have strong expectations about the anticipated effect size. However, previous research using the same task found medium-to-large effect sizes (i.e., *d*_z_ > 0.60) on half cycle responses for manipulations of fluency, also included here, and inversion, a manipulation of stimulus shape. Therefore, we conservatively assumed a medium-sized effect size (*d*_z_ = 0.50). An a-priori power analysis revealed that we needed at least 33 participants to detect such effect sizes with 80% power. Unfortunately, after testing the preregistered sample of *N* = 33, we discovered an undetected technical issue that resulted in bad data quality for 11 participants (i.e., > 10% of the electrodes requiring interpolation). Although we had not preregistered to compensate for excluded participants, the current study was conducted together with another study where we had preregistered to add 3 more participants if the sample size dropped below *N* = 30, until the sample size after exclusions was ≥ 30 (Cracco et al., 2022; preregistration: https://aspredicted.org/1P9_PNW). In that study, 9 participants had to be excluded and hence 6 participants were compensated. Given the large number of exclusions also in the current study, we decided to again combine both tasks for these 6 participants. As a result, the eventual sample size was *N* = 28 (9 male, 19 female, *M*_age_ = 23.14, range_age_ = 18–33). The experimental task procedures were conducted in accordance with the ethical protocol of the Faculty of Psychological and Educational Sciences at Ghent University.

### Task, Stimuli, and Procedure

Participants were seated in a Faraday cage, approximately 80-100 cm from a 24-inch computer monitor with a 60 Hz refresh rate. The experiment was programmed in PsychoPy (Peirce et al., 2019) and was based on Experiments 2-3 of Cracco et al. (2022). That is, participants watched videos of 4 identical agents arranged symmetrically around a fixation cross, synchronously performing the same movements (Figures 1-2). Note that four agents were shown instead of one to avoid overlap between the fixation cross and the moving stimuli. To control eye gaze and attention, participants were asked to focus on this fixation cross and to press the space bar every time it briefly (∼ 400 ms) turned red. Movements were presented as a repeating sequence of 12 images, rendered on the screen at a rate of 7.5 Hz. All sequences had the same symmetrical structure, with the second half of the sequence mirroring the first half. Previous research has shown that this elicits three brain responses: a brain response at 7.5 Hz (base rate), coupled to image presentation, a brain response at 1.25 Hz (half cycle rate), coupled to the symmetry point in the sequence, and a brain response at 0.625 Hz (full cycle rate), coupled to the repetition of the full image sequence (Cracco, Lee, et al., 2022).

**Figure 2.**
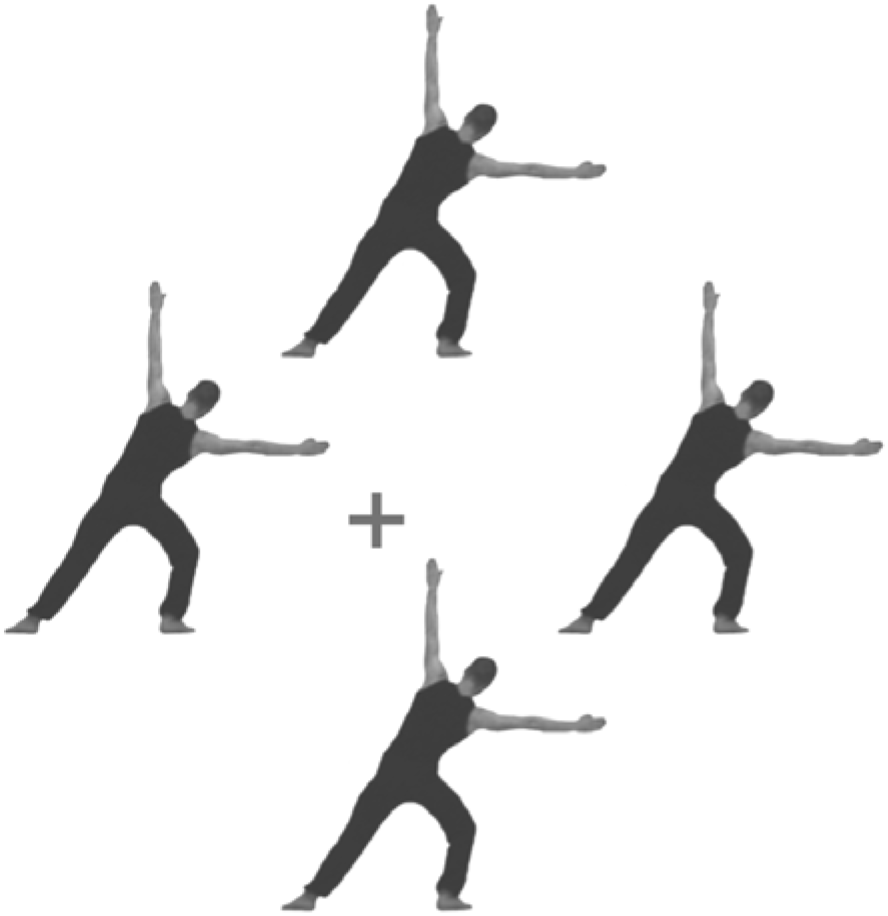
Example frame. Participants watched short videos showing four agents arranged symmetrically around a fixation cross, synchronously performing identical movements. Participants had to focus on the fixation cross and press the space bar each time it turned red. See Supplementary Video for an example video with non-pixelated human images.

Movement fluency was manipulated by changing the order of image presentation. In the fluent condition, images were ordered to depict agents performing a rhythmical dance movement, repeatedly moving their arms from left to right and back from right to left. In the non-fluent condition, images were rearranged to break this movement and to maximize visual displacement between successive body postures (Figure 1). Although both fluent and non-fluent sequences had the same symmetrical structure, this structure was more salient in the fluent condition because the half cycle point coincided with movement completion in this condition, signaling a change in movement direction (i.e. from left to right or from right to left). As a result, movement processing was captured at half cycle amplitudes. In contrast, image processing was captured at base rate and full cycle amplitudes (Cracco, Lee, et al., 2022).

In addition to movement fluency (fluent or non-fluent), we also manipulated stimulus animacy (human or corkscrew) and objecthood (defined shape or pixelated), resulting in 8 experimental conditions presented in random order in separate videos, with 3 repetitions per condition. Each video lasted for 60 sequence repetitions (i.e., 96 s), including a 3.2 s fade in and fade out to minimize eye blinking. Pixelated versions of the human and corkscrew images were created by degrading the pixel size with a 50×50 pixel size. All stimuli were matched for size as well as for contrast and luminance using the SHINE toolbox in Matlab 2020a. Nevertheless, it is not possible to fully match different stimuli for their apparent movement paths because the movement is fully determined by the shape and position of the stimulus. To compare the saliency of visual change between human, corkscrew, and pixelated image sequences, we therefore conducted a visual change analysis comparing the average pixel change between adjacent images of the different sequences (e.g., Orlandi et al., 2020; Table 1). This revealed that human and corkscrew stimuli were relatively matched for the saliency of apparent motion, differing primarily in their static, but not so much in their dynamic visual features. Instead, contrast changes between successive images were especially prominent for pixelated vs. normal sequences, and to a lesser extent non-fluent vs. fluent sequences. As a result, stronger brain responses for these sequences could be explained by increased stimulus flicker.

**Table 1.** Average Pixel Change Between Adjacent Images

|  | Normal |  | Pixelated |  |
| --- | --- | --- | --- | --- |
|  | Human | Corkscrew | Human | Corkscrew |
| Fluent | 1.35 | 1.39 | 6.08 | 5.61 |
| Non-Fluent | 1.87 | 1.62 | 8.46 | 6.67 |

### EEG Recording and Preprocessing

The EEG signal was recorded from 64 Ag/AgCI (active) electrodes using an ActiCHamp amplifier and BrainVisionRecorder software (version 1.21.0402, Brain Products, Gilching, Germany) at a sampling rate of 1000 Hz. Electrodes were positioned according to the 10%-system, except for two electrodes (TP9 and TP10), which were placed at OI1h and OI2h according to the 5%-system to have better coverage of posterior scalp sites. Fpz was used as ground electrode and Fz as online reference. Horizontal eye movements were recorded with two electrodes embedded in the EEG cap (FT9 and FT10) and vertical eye movements with two additional bipolar AG/AgCI sintered ring electrodes placed above and below the left eye.

Letswave 6 (www.letswave.org) was used for offline processing of the data. First, a fourth-order Butterworth bandpass filter with cut-off values 0.1 Hz and 100 Hz was applied. Next, the data were segmented according to the experimental conditions and ocular artefacts were removed using independent component analysis (ICA; RUNICA algorithm, square mixing matrix). After ICA, faulty or excessively noisy electrodes (2.8% on average, never more than 10%) were interpolated from the 3 closest neighbors and the data were re-referenced with respect to the average signal across electrodes. Finally, the fade in and out for each trial were cropped from the re-referenced epochs and averaged per condition before transforming them to normalized (divided by N/2) amplitudes (μV) in the frequency domain using a Fast Fourier Transform.

### Data Analysis

Brain responses at base rate (7.5 Hz), half cycle (1.25 Hz), and full cycle (0.625 Hz) were calculated as in Cracco et al. (2022). Amplitudes at each frequency bin were first baseline-corrected by subtracting the signal from the 10 neighboring frequency bins on each side (excluding directly adjacent bins) and the first 10 relevant harmonics of each response were then summed, excluding those harmonics that overlapped with the harmonics of a higher frequency response (consistent with the guidelines of Retter et al., 2021). The base rate harmonics included 7.5 Hz, 15 Hz, 22.5 Hz, 30 Hz, 37.5 Hz, 45 Hz, 52.5 Hz, 60 Hz, 67.5 Hz, and 75 Hz. The half cycle harmonics included 1.25 Hz, 2.50 Hz, 3.75 Hz, 5.00 Hz, 6.25 Hz, 8.75 Hz, 10 Hz, 11.25 Hz, 12.50 Hz, and 13.75 Hz. Finally, the full cycle harmonics included 0.625 Hz, 1.875 Hz, 3.125 Hz, 4.375 Hz, 5.625 Hz, 6.875 Hz, 8.125 Hz, 9.375 Hz, 10.625 Hz, and 11.875 Hz.

As in Cracco et al. (2022) we initially defined four electrode clusters to include in the analysis: a left posterior cluster (PO3, PO7, O1), a middle posterior cluster (Oz, OI1h, OI2h), a right posterior cluster (PO4, PO8, O2), and a middle frontocentral cluster (FCz, FC1, FC2). However, we decided to deviate from this preregistered approach because a collapsed localizer revealed (i) that there was no clear frontocentral activation in the current study and (ii) that the middle posterior cluster was not well captured by the preregistered electrodes (Figure 3)^1^. Accordingly, we did not include a frontocentral cluster in the analyses and used Oz, POz, and Pz to capture the middle posterior cluster rather than Oz, OI1h, and OI2h.

**Figure 3.**
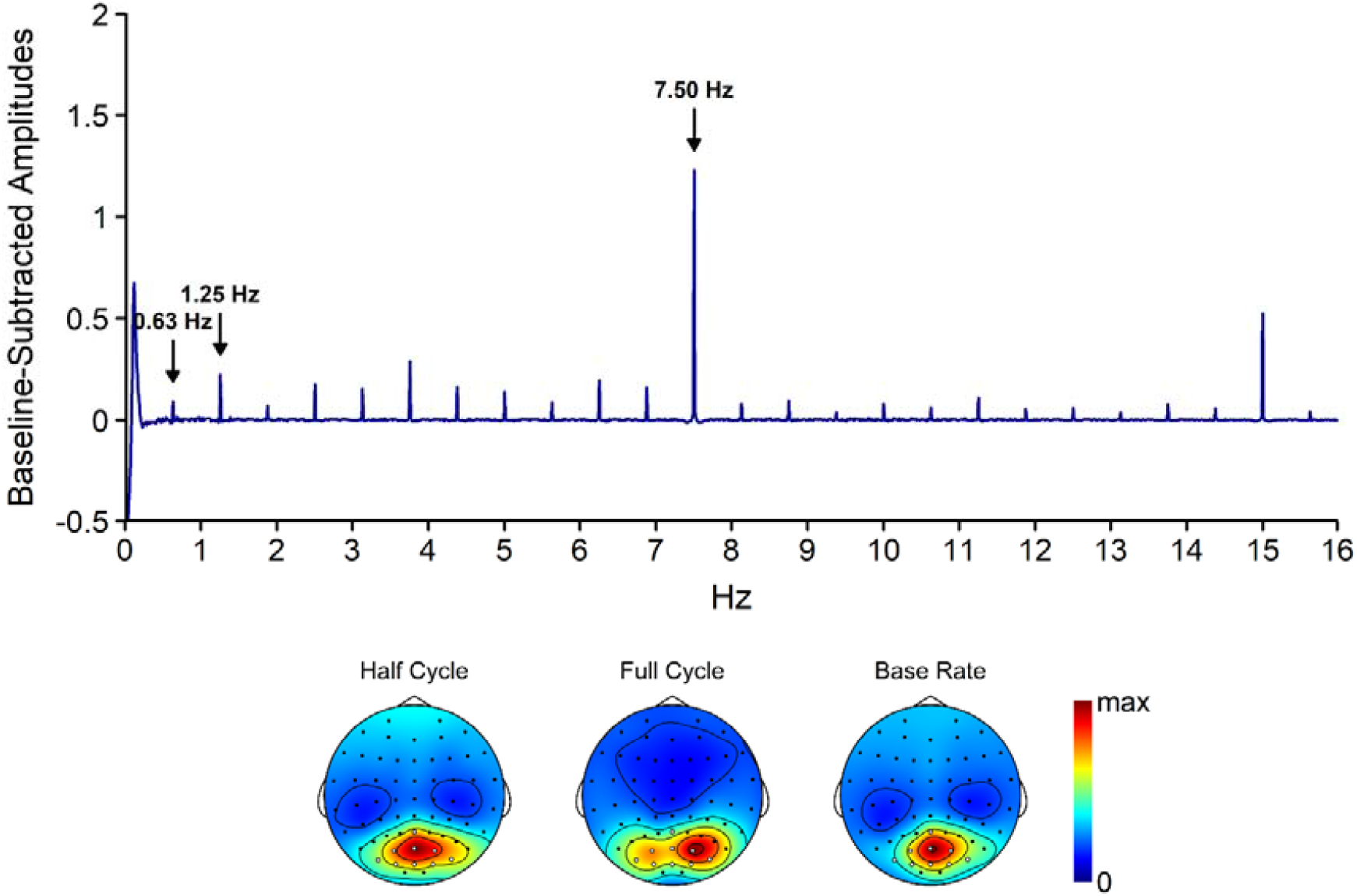
Collapsed spectrum plot and topographies of baseline-subtracted amplitudes. The spectrum plot shows the signal across the electrodes included into the analysis. The same electrodes are marked in white on the topographies. Topographies are scaled from 0 to the maximum value across the entire scalp for the respective response (i.e., 2.00 μV for the half cycle response, 1.38 μV for the full cycle response, and 3.35 μV for the base rate response). Separate topographies for the different conditions are reported in Supplemental Material.

Using these electrodes, we conducted a cluster (left, middle, or right posterior) x fluency (non-fluent or fluent) x pixelation (pixelated or normal) x animacy (corkscrew or human) Bayesian repeated measures ANOVA separately for half cycle, full cycle, and base rate amplitudes (Rouder et al., 2012). Note that we had initially planned to do a frequentist ANOVA but decided to instead conduct a more conservative Bayesian ANOVA to deal with the heightened false positive rate due to the large number of statistical tests. In addition, to simplify the results and because we were not interested in cluster effects, we added the main effect of cluster to the null model and did not fit interactions with cluster. Note, however, that the results did not change if we did fit these interactions, as shown in Supplementary Material. The Bayesian ANOVA was performed in JASP (JASP Team, 2022, v0.16.3) with default priors and using a model averaging approach that compared all models with the effect of interest to matched models without the effect of interest, excluding higher-order interactions (Wagenmakers et al., 2018). This yields a Bayes Factor (BF) for each effect, which represents the likelihood of the data under models containing vs. not containing the corresponding effect. A BF > 3 is typically considered reliable evidence for the presence of an effect, whereas a BF < 3 is typically considered reliable evidence against the presence of an effect (Jeffreys, 1961). If the Bayesian ANOVA revealed evidence for an effect with more than 2 levels, we followed up on this effect using Bayesian paired t-tests with default JASP priors. In addition to the BF, we also report η^p2^and Cohen’s *d*_z_ and to further quantify the size of each effect (Lakens, 2013).

## Results

### Half Cycle Rate

The brain response at half cycle rate is coupled to the symmetry point in the sequences. In fluent sequences, this corresponds to the completion of a body movement. Confirming that half cycle responses captured movement processing, the analysis of the half cycle response (Figure 3) revealed a main effect of fluency, BF_10_ = 10.54, d_z_ = 0.63, with stronger responses for fluent than for non-fluent sequences. In addition, there was a main effect of pixelation, BF_10_ = 1.45×10^7^, *d*_z_ = 1.71, with stronger responses for normal than for pixelated stimuli, and a main effect of animacy, BF_10_ = 109.52, *d*_z_ = 0.90, with stronger responses for corkscrews than for humans. The main effect of animacy was further qualified by a pixelation x animacy interaction, BF_10_ = 4.66, *d*_z_ = 0.47, indicating that the animacy effect was reliable for normal, BF_10_ = 1.31×10^3^, *d*_z_ = 0.97, but not for pixelated images, BF_10_ = 0.34, *d*_z_ = 0.20. No evidence was found for any of the other interaction effects, 0.23 ≤ BF_10_ ≤ 1.32.

### Full Cycle Rate

The brain response at full cycle rate is coupled to the completion of the full image sequence. In line with the half cycle analysis, the full cycle analysis (Figure 4) revealed main effects of fluency, BF_10_ = 1.38×10^5^, *d*_z_ = 1.38, pixelation, BF_10_ = 10.97, *d*_z_ = 0.64, and animacy, BF_10_ = 6.59×10^3^, *d*_z_ = 1.11. However, whereas the pixelation effect mirrored the half cycle results (normal > pixelated), the fluency (non-fluent > fluent) and animacy effects (human > corkscrew) were opposite. No evidence was found for any of the other effects, 0.44 ≤ BF_10_ ≤ 1.11.

**Figure 4.**
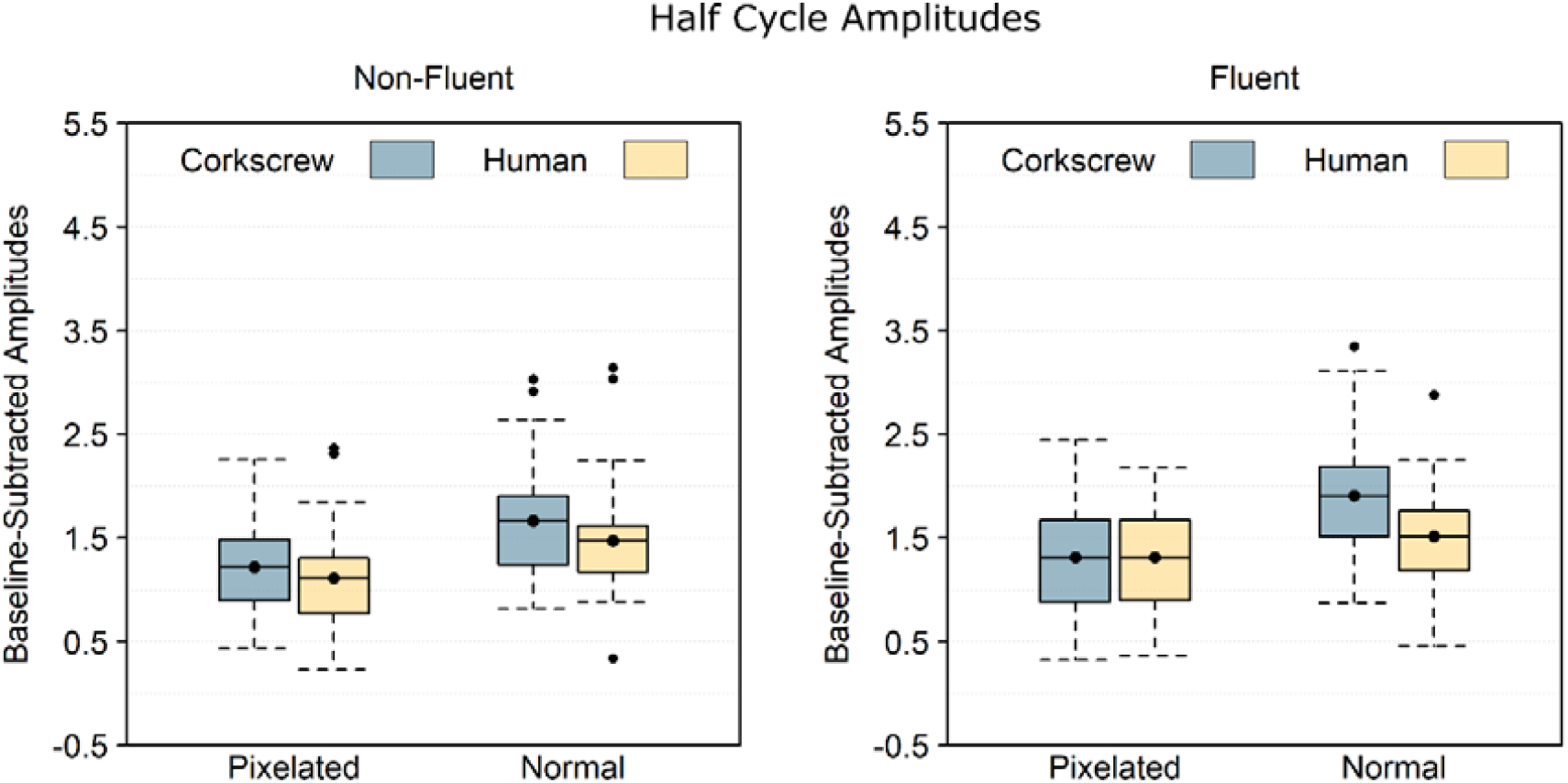
Baseline-subtracted amplitudes at half cycle rate (1.25 Hz) and its harmonics. Boxplots show the mean instead of the median to better match the statistical analysis. Note that 0 is the baseline and that values below 0 necessarily reflect noise. The black dots are data points exceeding the first or third quartile by more than 1.5 times the interquartile range.

### Base Rate

The brain response at base rate is coupled to the presentation of the individual images. The base rate analysis (Figure 5) revealed reliable main effects of fluency, BF_10_ = 3.28, *d*_z_ = 0.52, and pixelation, BF_10_ = 60.29, *d*_z_ = 0.76, but not animacy, BF_10_ = 2.90, *d*_z_ = 0.45. Interestingly, while the fluency effect (non-fluent > fluent) mirrored the full cycle response, the pixelation effect was opposite (pixelated > normal). In addition to these main effects, there was also a pixelation x animacy interaction, BF_10_ = 6.11, *d*_z_ = 0.54, indicating that that the pixelation effect was reliable for human, BF_10_ = 1.06×10^3^, *d*_z_ = 0.97, but not for corkscrew sequences, BF_10_ = 0.83, *d*_z_ = 0.34. No evidence was found for any of the other effects, 0.41 ≤ BF_10_ ≤ 1.68.

**Figure 5.**
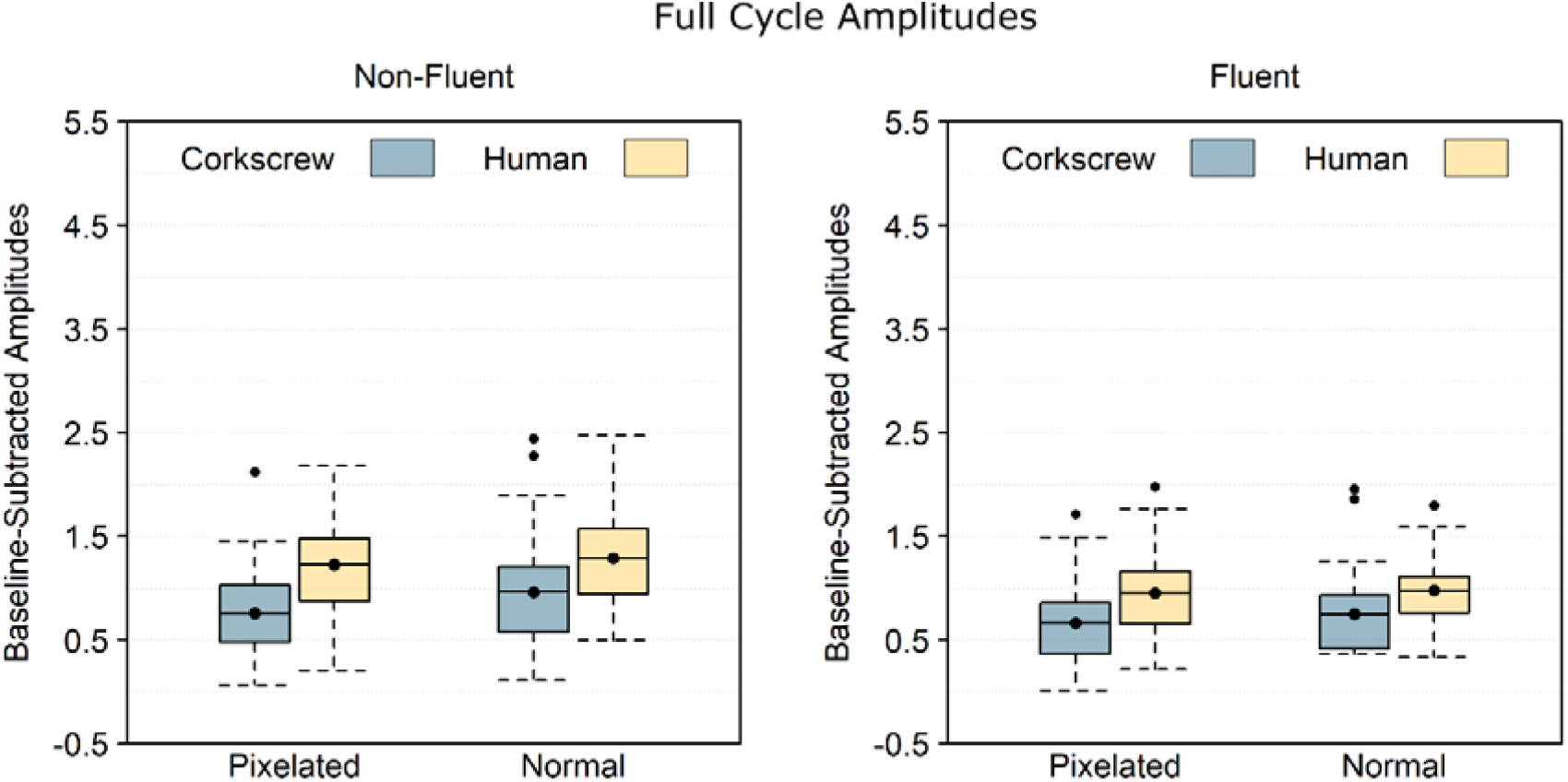
Baseline-subtracted amplitudes at full cycle rate (0.625 Hz) and its harmonics. Boxplots show the mean instead of the median to better match the statistical analysis. Note that 0 is the baseline and that values below 0 necessarily reflect noise. The black dots are data points exceeding the first or third quartile by more than 1.5 times the interquartile range.

**Figure 6.**
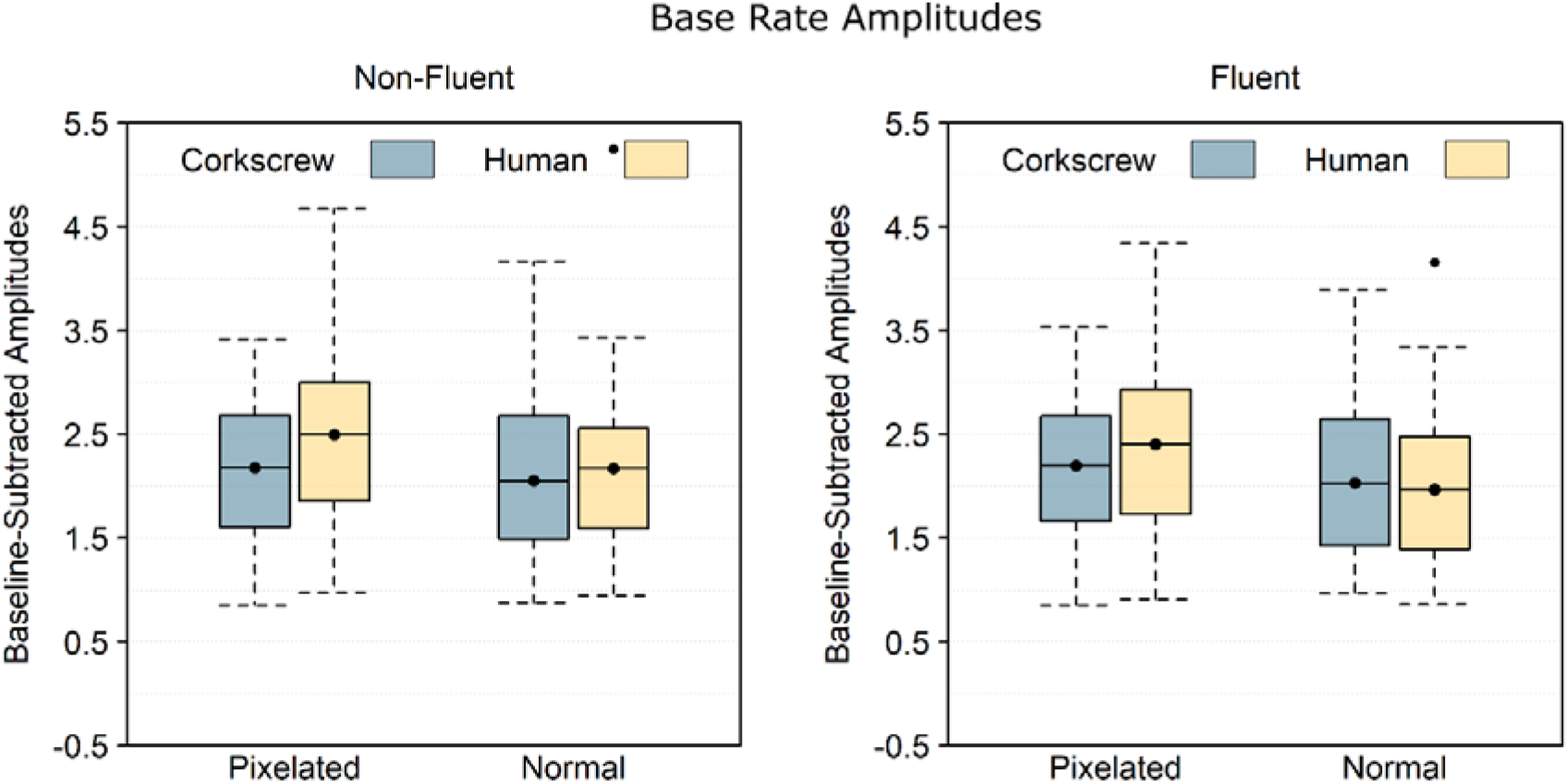
Baseline-subtracted amplitudes at base rate (7.5 Hz) and its harmonics. Boxplots show the mean instead of the median to better match the statistical analysis. Note that 0 is the baseline and that values below 0 necessarily reflect noise. The black dots are data points exceeding the first or third quartile by more than 1.5 times the interquartile range.

## Discussion

This study investigated the influence of objecthood and animacy on movement processing in the form pathway. To this end, we showed normal and pixelated image sequences of human and corkscrew stimuli producing fluent and non-fluent apparent movements. These movements occurred at half the rate of sequence presentation, eliciting corresponding brain responses that can be used as a measure of movement processing (Cracco, Lee, et al., 2022). Supporting the hypothesis that objecthood would influence movement processing, we found weaker half cycle responses for pixelated than for non-pixelated stimuli. In contrast, we found no evidence for human-specificity, with stronger as opposed to weaker responses for corkscrew than for the human stimuli. Finally, although we found overall stronger half cycle responses for fluent than for non-fluent sequences (Cracco, Lee, et al., 2022), neither the objecthood nor the animacy effect depended on movement fluency.

The absence of interactions with movement fluency indicates that neither the effect of objecthood, nor the effect of animacy was specific to movement processing. Indeed, scrambling the order of images in an apparent movement sequence is well-known to disrupt movement processing (Cracco, Lee, et al., 2022; Orgs et al., 2016), as it makes it more difficult to integrate images over time into a coherent movement percept (Lange & Lappe, 2006, 2007). Importantly, however, binding postures into meaningful chunks is not necessarily unique to movement processing and postures could also be structured according to different criteria. For instance, sequences were symmetrical around the half cycle point in our task. As a result, posture integration could have also been driven by symmetry, a well-known principle of perceptual organization (Wagemans et al., 2012). Applied to objecthood, this would mean that binding stimuli into chunks is easier when they have an easily recognizable, familiar shape (main effect of pixelation) and that this effect is relevant for movement processing (main effect of movement fluency), but not specific to movement processing (no interaction).

Interestingly, this interpretation is at odds with a previous fMRI study that did find a movement-specific effect of stimulus pixelation (Orgs et al., 2016). However, movement specificity was limited to motor areas. Brain activity in the current study, on the other hand, was restricted to posterior brain areas. It is, therefore, possible that movement-specific effects of objecthood exist only in the motor cortex. If so, this would suggest that they are driven by motor simulation, the process of mentally simulating other people’s actions in one’s own motor system (Jeannerod, 2001). Although frontal brain activity indicative of motor simulation was observed in three previous experiments using the same paradigm (Cracco, Lee, et al., 2022), the same frontal activation cluster was not found here. While speculative, this is potentially the result of mixing sequences with human and non-human agents. Indeed, motor simulation is known to be reduced or absent for non-human agents (Cracco et al., 2018; Press, 2011) and mixing human with non-human sequences may therefore have led to a more perceptual processing style as opposed to previous studies (Cracco, Lee, et al., 2022; Orgs et al., 2016).

In contrast to objecthood, stimulus animacy had the opposite effect as predicted, namely stronger responses for the corkscrew than for the human agent. A likely explanation for this finding is that the movements made by the corkscrew stimulus were more salient. This could have had several reasons. First, it is possible that the corkscrew’s movements were more salient because it is unusual to see an inanimate object move independently. Second, consistent with evidence that apparent movement perception depends on the rate at which the images are presented (Grosjean et al., 2007; Shiffrar & Freyd, 1990), the corkscrew’s movements may have been more salient because the presentation rate was more suited for the corkscrew than for the human stimulus. Nevertheless, regardless of the explanation, it is clear that, at the very least, animacy is not the primary force driving movement processing in the form pathway. These findings are consistent with previous work showing that even simple geometrical shapes can elicit vivid illusions of social interaction (Heider & Simmel, 1944; Ramsey & Hamilton, 2010), suggesting that the brain also attributes animacy to non-human stimuli.

In addition to brain responses at half cycle rate, we also measured two other responses: brain responses at full cycle rate and brain responses at base rate. In a previous study, we interpreted full cycle responses as reflecting the processing of body posture sequences and found that they were stronger when the individual postures became more salient, such as in non-fluent sequences (Cracco, Lee, et al., 2022). The current results not only replicate this finding but further bolster it by showing that full cycle responses were also stronger for human than for blurred and non-human postures. These results are consistent with previous research showing that there are dedicated brain areas for processing human body images, such as the fusiform (Schwarzlose et al., 2005) and extrastriate body area (Downing et al., 2006), and suggest that the full cycle response, unlike the half cycle response, is specific to human body stimuli.

The base rate response, because it is coupled to image presentation, was previously interpreted to capture (body) image processing regardless of posture (Cracco, Lee, et al., 2022). However, for the same reason, it is also likely to capture changes in contrast from image-to-image. Supporting the latter view, the base rate response tended to be stronger for sequences where visual change was larger (Table 1). Indeed, both the base rate and visual change analyses found the largest differences between pixelated and normal sequences, then between non-fluent and fluent sequences, and finally between human and corkscrew sequences. It therefore seems likely that base rate responses primarily captured low-level visual processing. This same reasoning also implies that low-level visual processing was likely not captured by half and full cycle responses, as they were both reduced for pixelated stimuli.

In sum, the current study shows that movement processing in the form pathway, like movement processing in the motion pathway, requires a familiar shape, but not necessarily a human shape (Jastorff et al., 2006, 2009). This was true even though human-specificity did appear at the level of posture processing and likely indicates that movement perception in the form pathway relies on domain-general mechanisms that are used not only for movement processing but also to process other sorts of structured sequences, as supported by the lack of interactions with movement fluency in the current study. Overall, our findings thus suggest that there is at least a partial dissociation between body and movement processing in the form pathway.

## Supplementary Material

**Half Cycle Results With Cluster**

**Table S1.** Half Cycle Results Table

| Effect | BF<sub>10</sub> | Pattern |
| --- | --- | --- |
| Cluster | 67.86 | M > L<br>R > L<br>M ? R |
| Fluency | 11.85 | F > NF |
| Agent | 117.79 | C > H |
| Pixelation | 1.34x10<sup>7</sup> | N > P |
| Cluster x Fluency | 0.07 | NA |
| Cluster x Agent | 3.03 | L: C > H<br>M: C ? H<br>R: C > H |
| Cluster x Pixelation | 0.17 | NA |
| Fluency x Agent | 0.36 | NA |
| Fluency x Pixelation | 0.30 | NA |
| Agent x Pixelation | 5.47 | P: C = H<br>N: C > H |
| Cluster x Fluency x Agent | 0.20 | NA |
| Cluster x Fluency x Pixelation | 0.19 | NA |
| Cluster x Agent x Pixelation | 2.30 | NA |
| Fluency x Agent x Pixelation | 0.57 | NA |
| Cluster x Fluency x Agent x Pixelation | 8.19x10<sup>-15</sup> | NA |

*Note.* BF<sub>10</sub> ≤ 0.33 is denoted with “=”, 0.33 < BF<sub>10</sub> < 3 with “?”, and BF<sub>10</sub> ≥ 3 with “>”. L:
left, M: middle, R: right, NF: non-fluent, F: fluent, C: corkscrew, H: human, P: pixelated, N: normal.

**Table S2.** Means and Standard Deviations of the Half Cycle Response

| **Left** | Non-Fluent |  | Fluent |  |
| --- | --- | --- | --- | --- |
|  | Pixelated | Normal | Pixelated | Normal |
| Cokscrex | $1.01 \pm 0.51$ | $1.64 \pm 0.74$ | $1.12 \pm 0.65$ | $1.80 \pm 0.81$ |
| Human | $0.98 \pm 0.69$ | $1.25 \pm 0.71$ | $1.13 \pm 0.62$ | $1.35 \pm 0.59$ |
| **Middle** | Non-Fluent |  | Fluent |  |
|  | Pixelated | Normal | Pixelated | Normal |
| Cokscrex | $1.37 \pm 0.67$ | $1.71 \pm 0.93$ | $1.47 \pm 0.83$ | $1.97 \pm 1.14$ |
| Human | $1.29 \pm 0.81$ | $1.73 \pm 1.04$ | $1.54 \pm 0.92$ | $1.67 \pm 0.97$ |
| **Right** | Non-Fluent |  | Fluent |  |
|  | Pixelated | Normal | Pixelated | Normal |
| Cokscrex | $1.27 \pm 0.58$ | $1.64 \pm 0.76$ | $1.34 \pm 0.73$ | $1.95 \pm 0.92$ |
| Human | $1.07 \pm 0.72$ | $1.44 \pm 0.76$ | $1.26 \pm 0.80$ | $1.52 \pm 0.72$ |

**Figure S1.**
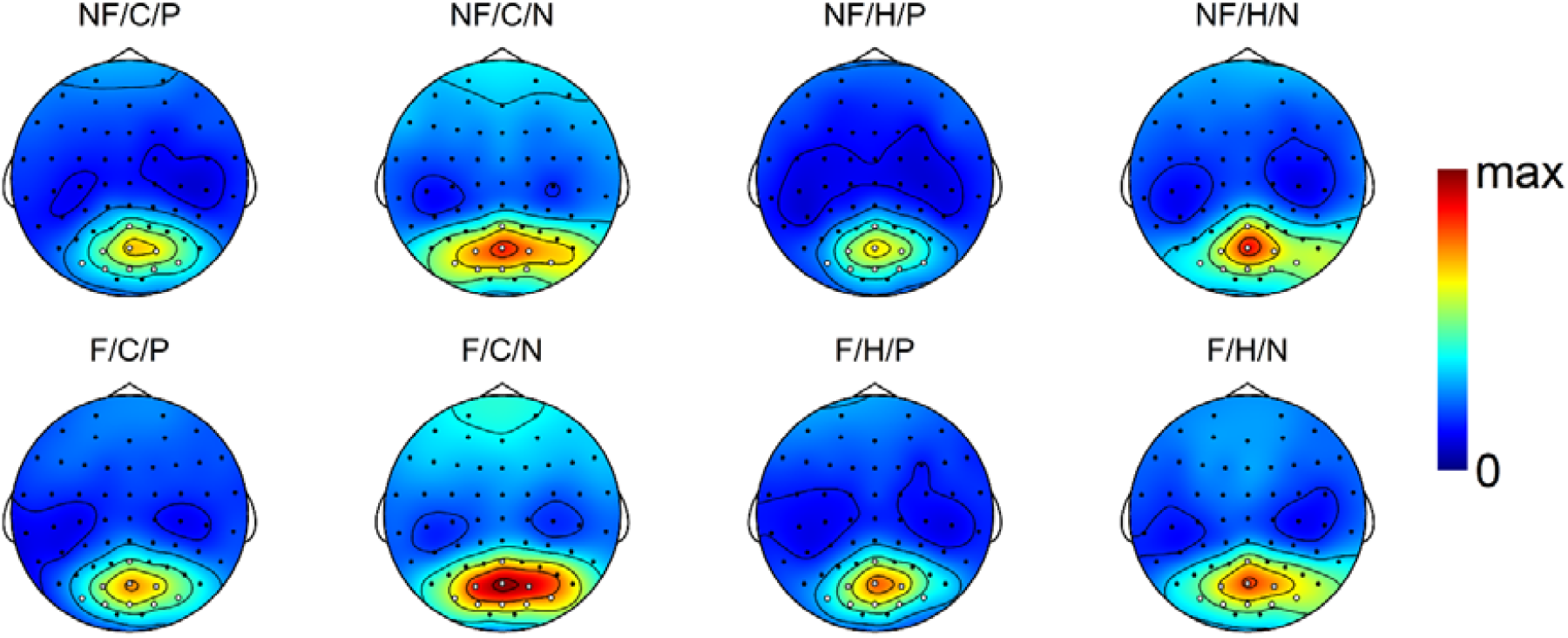
Condition-specific topographies for the half cycle amplitudes. All topographies are scaled from 0 to the maximum value across the scalp and across the different conditions (i.e., 2.50 μV). NF: non-fluent, F: fluent, C: corkscrew, H: human, P: pixelated, N: normal.

**Full Cycle Results With Cluster**

**Table S3.** Full Cycle Results Table

| Effect | BF<sub>10</sub> | Pattern |
| --- | --- | --- |
| Cluster | 194.80 | R > M<br>R > L<br>L ? M |
| Fluency | 1.32x10<sup>5</sup> | NF > F |
| Agent | 6.60x10<sup>3</sup> | H > C |
| Pixelation | 10.82 | N > P |
| Cluster x Fluency | 0.10 | NA |
| Cluster x Agent | 3.57x10<sup>-7</sup> | NA |
| Cluster x Pixelation | 3.60 | L: N > P<br>M: N = P<br>R: N ? P |
| Fluency x Agent | 1.14 | NA |
| Fluency x Pixelation | 0.75 | NA |
| Agent x Pixelation | 0.88 | NA |
| Cluster x Fluency x Agent | 9.16x10<sup>-4</sup> | NA |
| Cluster x Fluency x Pixelation | 0.11 | NA |
| Cluster x Agent x Pixelation | 6.47 | Unclear |
| Fluency x Agent x Pixelation | 0.19 | NA |
| Cluster x Fluency x Agent x Pixelation | 2.00x10<sup>-3</sup> | NA |

*Note.* BF<sub>10</sub> ≤ 0.33 is denoted with “=”, 0.33 < BF<sub>10</sub> < 3 with “?”, and BF<sub>10</sub> ≥ 3 with “>”. L:
left, M: middle, R: right, NF: non-fluent, F: fluent, C: corkscrew, H: human, P: pixelated, N: normal. The pattern underlying the cluster x agent x pixelation interaction suggested the presence of an agent x pixelation interaction for the middle cluster but not for the other two clusters. However, this pattern could not be supported by post-hoc Bayesian paired *t*-tests.

**Table S4.** Means and Standard Deviations of the Full Cycle Response

| **Left** | Non-Fluent |  | Fluent |  |
| --- | --- | --- | --- | --- |
|  | Pixelated | Normal | Pixelated | Normal |
| Cokscrex | $0.71 \pm 0.67$ | $0.93 \pm 0.78$ | $0.63 \pm 0.61$ | $0.77 \pm 0.69$ |
| Human | $1.14 \pm 0.67$ | $1.36 \pm 0.77$ | $0.90 \pm 0.55$ | $1.01 \pm 0.60$ |
| **Middle** | Non-Fluent |  | Fluent |  |
|  | Pixelated | Normal | Pixelated | Normal |
| Cokscrex | $0.65 \pm 0.56$ | $0.79 \pm 0.67$ | $0.57 \pm 0.45$ | $0.57 \pm 0.44$ |
| Human | $1.12 \pm 0.70$ | $1.06 \pm 0.81$ | $0.86 \pm 0.64$ | $0.75 \pm 0.58$ |
| **Right** | Non-Fluent |  | Fluent |  |
|  | Pixelated | Normal | Pixelated | Normal |
| Cokscrex | $0.91 \pm 0.77$ | $1.17 \pm 0.94$ | $0.79 \pm 0.67$ | $0.91 \pm 0.77$ |
| Human | $1.43 \pm 0.72$ | $1.45 \pm 0.85$ | $1.09 \pm 0.65$ | $1.17 \pm 0.62$ |

**Figure S2.**
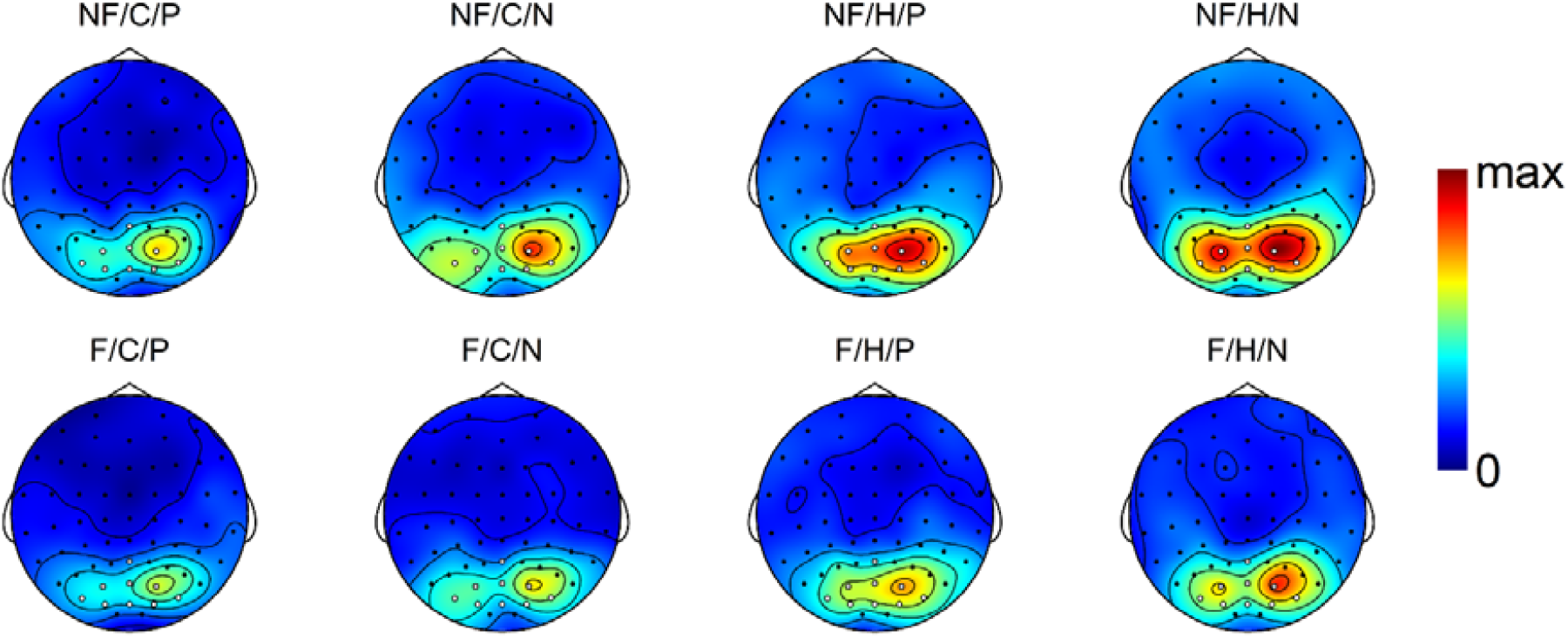
Condition-specific topographies for the half cycle amplitudes. All topographies are scaled from 0 to the maximum value across the scalp and across the different conditions (i.e., 1.77 μV). NF: non-fluent, F: fluent, C: corkscrew, H: human, P: pixelated, N: normal.

**Base Rate Results With Cluster**

**Table S5.** Base Rate Results Table

| Effect | BF<sub>10</sub> | Pattern |
| --- | --- | --- |
| Cluster | 3.45x10<sup>4</sup> | M > L<br>M > R<br>R > L |
| Fluency | 4.21 | NF > F |
| Agent | 3.10 | H > P |
| Pixelation | 60.29 | P > N |
| Cluster x Fluency | 0.20 | NA |
| Cluster x Agent | 1.25 | NA |
| Cluster x Pixelation | 29.99 | L: P ? N<br>M: P > N<br>R: P > N |
| Fluency x Agent | 1.45 | NA |
| Fluency x Pixelation | 1.53 | NA |
| Agent x Pixelation | 6.72 | C: P ? N<br>H: P > N |
| Cluster x Fluency x Agent | 0.18 | NA |
| Cluster x Fluency x Pixelation | 0.14 | NA |
| Cluster x Agent x Pixelation | 3.29 | L: Agent x Pixelation ?<br>M: Agent x Pixelation +<br>R: Agent x Pixelation ? |
| Fluency x Agent x Pixelation | 0.40 | NA |
| Cluster x Fluency x Agent x Pixelation | 13.78 | Overcomplex |

*Note.* BF<sub>10</sub> ≤ 0.33 is denoted with “=”, 0.33 < BF<sub>10</sub> < 3 with “?”, and BF<sub>10</sub> ≥ 3 with “>”. L:
left, M: middle, R: right, NF: non-fluent, F: fluent, C: corkscrew, H: human, P: pixelated, N:
normal. The 4-way interaction was considered too complex to interpret and was not further analyzed.

**Table S6.** Means and Standard Deviations of the Base Rate Response

| Left | Non-Fluent |  | Fluent |  |
| --- | --- | --- | --- | --- |
|  | Pixelated | Normal | Pixelated | Normal |
| Cokscrex | $1.63 \pm 0.86$ | $1.71 \pm 0.96$ | $1.70 \pm 0.89$ | $1.63 \pm 0.90$ |
| Human | $1.99 \pm 1.01$ | $1.83 \pm 0.93$ | $1.82 \pm 0.92$ | $1.59 \pm 0.82$ |
| Middle | Non-Fluent |  | Fluent |  |
|  | Pixelated | Normal | Pixelated | Normal |
| Cokscrex | $2.64 \pm 1.25$ | $2.42 \pm 1.20$ | $2.66 \pm 1.25$ | $2.41 \pm 1.25$ |
| Human | $3.13 \pm 1.47$ | $2.52 \pm 1.25$ | $3.03 \pm 1.45$ | $2.39 \pm 1.16$ |
| Right | Non-Fluent |  | Fluent |  |
|  | Pixelated | Normal | Pixelated | Normal |
| Cokscrex | $2.25 \pm 1.27$ | $2.03 \pm 1.24$ | $2.23 \pm 1.25$ | $2.05 \pm 1.26$ |
| Human | $2.37 \pm 1.25$ | $2.16 \pm 1.27$ | $2.37 \pm 1.41$ | $1.93 \pm 1.16$ |

**Figure S3.**
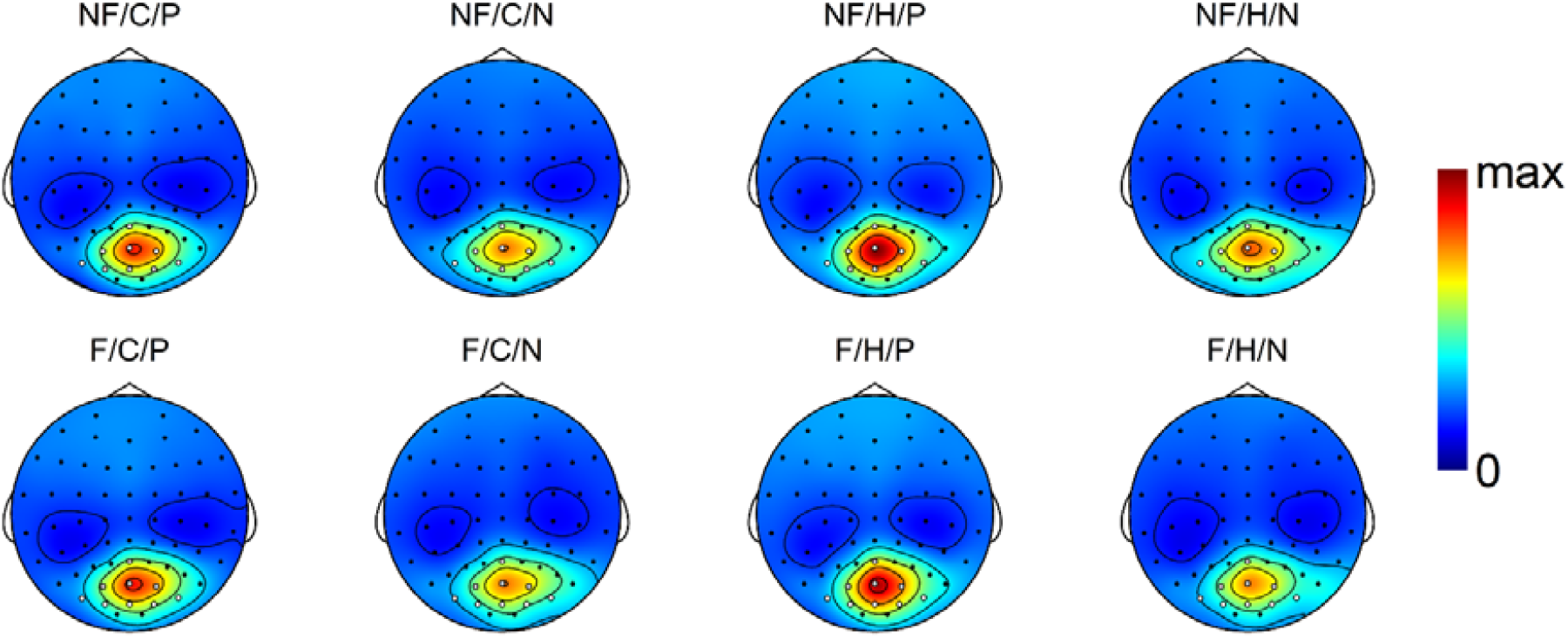
Condition-specific topographies for the half cycle amplitudes. All topographies are scaled from 0 to the maximum value across the scalp and across the different conditions (i.e., 3.99 μV). NF: non-fluent, F: fluent, C: corkscrew, H: human, P: pixelated, N: normal.

## Footnotes

1 Note that this is likely because this study uses a different electrode setup that does not include all electrodes included in the middle posterior cluster of the original study of Cracco, Lee, et al. (2022).

